# Temperament profile and its association with the vulnerability to smartphone addiction of medical students in Indonesia

**DOI:** 10.1101/536474

**Authors:** Enjeline Hanafi, Kristiana Siste, Tjhin Wiguna, Irmia Kusumadewi, Martina Wiwie Nasrun

## Abstract

Temperament profiles of an individual with high novelty seeking and low harm avoidance have been reported to be related to substance addiction, but smartphone addiction could be different from substance addiction. Medical students have high exposure to smartphone use. Screening their risk of smartphone addiction based on the temperament profile may help in deciding best prevention approach. This research aimed to examine the temperament profile and its association with vulnerability to smartphone addiction of medical students in Jakarta, Indonesia. The research was conducted with cross sectional design and simple random sampling. The Temperament and Character Inventory and the Smartphone Addiction Scale were used to measure desired outcomes. Logistic regression analysis was performed to identify the relationship between demographic factors, pattern of smartphone use, temperament type, and vulnerability to smartphone addiction. Of the 185 samples, most subjects have low novelty seeking, high reward dependence, and high harm avoidance. The average smartphone usage in a day was 7.94 hours (SD 3.92) with the initial age of smartphone usage was 7.58 years (SD 2.43). The respondents used smartphone for communication and accessing social media. High harm avoidance temperament was significantly associated with risk of smartphone addiction with OR 2.035; 95% CI 1.119 to 3.701). This study shows that smartphone addiction has similarities with other behavior addictions. Harm avoidance temperament is associated with the risk of smartphone addiction. Screening on risk of smartphone addiction based on temperament type should be done on medical students.

## Introduction

Smartphones, mobile phones that provide integrated communication and computation services, play an important role in people’s lives worldwide (1,2). According to a study by Pew Research Center, smartphone ownership strongly correlates with internet connectivity (3). The numbers of smartphone owner and internet user have increased drastically in the last couple of years. In developing countries, smartphone ownership has climbed from 21% in 2013 to 37% in 2015 (3). Approximately 21% of Indonesians own a smartphone in 2015 (3). Smartphone owners in Indonesia are more likely to be 18 to 34 years old, highly-educated, and have high income (3). The most popular activities on the smartphone are social networking through sites such as Facebook®, Twitter®, dan Path® (3).

The high number of smartphone ownership raises questions about its risks. In general, smartphone use is associated with motor vehicle accidents, academic problems, and health issues (4). More frequent use of smartphone increases the prevalence of sleep disorders, depression, and anxiety (5). Smartphone overuse will have a significant impact.

In a more severe condition, smartphone use can cause dependence or addiction. A study reports that 46% of smartphone users feel that they cannot live without smartphones (3). Another study found that, on average, university students checks their phones 60 times a day, with more than 4 hours of regular use. This fear of separation from smartphone has been termed nomophobia (6). When separation occurs, withdrawal-like restlessness and physiological symptoms emerge. Some report that they heard their phones ring or vibrate, when they actually do not. This phenomenon is not the same as compensation, as its objective is neither escape from problems or responsibilities, nor avoidance of negative emotion. Considering these aspects, when a person becomes addicted to smartphones, the symptoms of addiction and withdrawal can cause impairment in that person’s life.

Early detection is the first step in mitigating the consequences of smartphone addiction and should take individual risk factors into consideration. Temperament is one of such risk factors. Currently, there is no study exploring the association between temperament profile and smartphone addiction. Temperament is the emotional pattern of an individual, free from the influence of values and norms, and is associated with specific systems in the brain that regulates negative emotion and behavior. Cloninger divides temperament into three groups: *novelty seeking* (NS), *reward dependence* (RD), and *harm avoidance* (HA) (7). The temperament profile is important in understanding the reason behind someone’s thought, perception, and behavior. Ko et al. has conducted a study of temperament in behavioral addiction (8). They found that people with internet addiction has high NS, low RD, and high HA. However, such study has never been done on those with smartphone addiction. Therefore, further research is needed because risk factors for internet addiction may not apply in smartphone addiction.

According to Eley et al., Australian medical students tend to have high HA, similar to those with internet addiction (9,10). This is quite concerning if medical students in Jakarta have similar temperament profile, as students are required to keep up to date on medical information and they rely on convenient devices such as smartphones and other gadgets (11). Additionally, medical students are a group with long study period and high expectations (12). Smartphones that were initially used for academic purposes later become an escape. If this is neglected, there will be consequences on academic achievement, daily life, and their future (13). Before the students become addicted to smartphones, prevention must be done, for example, through screening, but screening is not feasible to be conducted on all students. Screening their risk of smartphone addiction based on the temperament profile may help in deciding appropriate approach in preventing or managing the condition. This study aims to identify the temperament profile of medical students in Jakarta and its association to vulnerability to smartphone addiction.

## Methods

This is a cross-sectional study, conducted in several medical faculties in Jakarta. Data collection was performed from January to April 2018.

### Participants

The sample for this study was all preclinical medical students in Jakarta who fulfil the inclusion and exclusion criteria. Sample selection was done by simple random sampling, randomized using the software for analysis.

The inclusion criteria are those who were preclinical medical students in the fifth semester, had completely filled out demographic form and both questionnaires, and had agreed to be a respondent by signing the informed consent. Subject was excluded if he/she was not fluent in Indonesia Language.

### Measurements

#### Modified SAS Indonesian version

Vulnerability to smartphone addiction was measured using SAS, a self-report scale. This instrument has 33 questions with a likert-scale response of 1 to 6. There are 6 subscales: cyberspace-oriented relationship, daily life disruption, primary needs, overuse, positive anticipation, and withdrawal (14). Cronbach’s alpha score for the whole scale is 0.967. This instrument is valid, as compared to similar instruments. SAS has been validated into Indonesian (15). SAS Indonesian version was modified into 21 questions with item-total correlation of 0.282 to 0.802 and Cronbach’s alpha of 0.890. SAS produces a numerical score, with a higher score signifying a higher vulnerability to smartphone addiction.

#### Modified TCI Indonesian version

To identify a patient’s temperament type, modified TCI Indonesian version was used (16). This self-report instrument has 23 questions on temperament (9 items on novelty seeking, 6 on reward dependence, and 8 on harm avoidance) and 16 questions on character (10 items on self-directedness and 6 on self-transcendence). This study utilized only the questions on temperament.

Temperament is measured using 23 items answered either ‘yes’ (2 points) or ‘no’ (1 point) according to respondent’s view of themselves. There is an answer key for the items. The output of this instrument is the sum of each temperament group. Higher score on each item means higher scores on harm avoidance, novelty seeking, and reward dependence. The low and high cut-off point for each group is determined by the average of all subjects.

### Ethical approval

This study was cleared by the local Ethics Committee. Study subjects were asked to sign informed consent.

### Statistical Analysis

Data was collected and then analyzed statistically. Data was processed using *Statistical Analysis Software Package for Windows* (SPSS®, IBM, USA) version 25. Demographic factors were summarized into proportion for nominal variables, mean and standard deviation for numeric variables. Logistic regression analysis was performed to identify the relationship between demographic factors, pattern of smartphone use, temperament type, and vulnerability to smartphone addiction.

## Results

This cross-sectional study was conducted on 300 medical students in three medical faculties on their fourth or fifth semester. Only 258 forms fulfilled the study criteria. Sample was then randomly selected to obtain 185 samples.

### Demographic Characteristics of Study Subjects

The mean age of study subjects is 20.39 years (standard deviation 1.14). Most are females (66.5%) and none have married (Table 1).

**Table 1.** Demographic Characteristics of Study Subjects.

|  | Mean (SD) | 95% CI |  | n | % |
| --- | --- | --- | --- | --- | --- |
|  |  | Lower limit | Upper limit |  |  |
| **Age (years)** | 20.39 (1.14) | 20.23 | 20.56 |  |  |
| **Marital status** |  |  |  |  |  |
| Married |  |  |  | 0 | 0 |
| Not married |  |  |  | 185 | 100 |
| **Gender** |  |  |  |  |  |
| Male |  |  |  | 62 | 33.5 |
| Female |  |  |  | 123 | 66.5 |

SD, standard deviation; CI, confidence interval

### Smartphone Usage Characteristics

Study subjects mostly used their smartphones for communication (41.1%) and to access social media (29.7%). The mean duration of smartphone use each day is 7.83 hours (SD 4.03), with 68.6% of subjects using their smartphones for 6 hour or more. Mean age at first use of smartphone is 7.62 (SD 2.60; Table 2).

**Table 2.** Smartphone Usage Characteristics.

|  | Mean (SD) | 95% CI |  | n | % |
| --- | --- | --- | --- | --- | --- |
|  |  | Lower limit | Upper limit |  |  |
| Age at first use (years) | 7.62 (2.60) | 7.24 | 8.00 |  |  |
| Duration of daily use (hours) | 7.83 (4.03) | 7.24 | 8.41 |  |  |
| ≥ 6 hours |  |  |  | 127 | 68.6 |
| < 6 hours |  |  |  | 58 | 31.4 |
| Activity done using smartphone |  |  |  |  |  |
| Communication |  |  |  | 76 | 41.1 |
| Browsing |  |  |  | 26 | 14.1 |
| Games |  |  |  | 12 | 6.5 |
| Social media |  |  |  | 55 | 29.7 |
| Entertainment |  |  |  | 16 | 8.6 |

SD, standard deviation; CI, confidence interval.

### Smartphone Activities According to Gender

Males are more likely to use smartphones for communication (35.5%), social media (24.2%), and games (19.4%). On the other hand, females are more likely to use smartphones for communication (43.1%), social media (33.3%), and browsing (13.8%). There is a significant difference between males and females in the use of smartphones for games (X^2^ = 21,69, p < 0.001, Table 3).

**Table 3.** Smartphone Activities in Males and Females.

| Activity | Gender |  | p value |
| --- | --- | --- | --- |
|  | Male | Female |  |
| Communication | 23 (37.1%) | 53 (43.1%) | 0.527 |
| *Browsing* | 10 (16.1%) | 16 (13.0%) | 0.655 |
| Games | 10 (16.1%) | 2 (1.6%) | <0.001* |
| Social media | 14 (22.6%) | 41 (33.3%) | 0.173 |
| Entertainment | 5 (8.1%) | 11 (8.9%) | 1.000 |

\* $p < 0.05$

### Temperament Profile of Study Subjects

Mean scores for temperament groups NS, RD, and HA are 11.56 (SD 1.89); 9.36 (SD 1.69), and 11.74 (SD 1.88), respectively. Those scoring above these mean scores are considered high in that specific temperament. More than 50% subjects have low NS, high RD, and high HA (Table 4).

**Table 4.** Temperament Profile of Study Subjects.

| Temperament | Mean (SD) | 95% CI |  | n | % |
| --- | --- | --- | --- | --- | --- |
|  |  | Lower limit | Upper limit |  |  |
| *Novelty Seeking* | 11.67 (1.80) | 11.41 | 11.93 |  |  |
| High |  |  |  | 89 | 48.1 |
| Low |  |  |  | 96 | 51.9 |
| *Reward Dependence* | 9.44 (1.66) | 9.20 | 9.68 |  |  |
| High |  |  |  | 98 | 53.0 |
| Low |  |  |  | 87 | 47.0 |
|  |  | Lower limit | Upper limit |  |  |
| *Harm Avoidance* | 11.65 (1.91) | 11.37 | 11.93 |  |  |
| High |  |  |  | 98 | 53.0 |
| Low |  |  |  | 87 | 47.0 |

SD, standard deviation; CI, confidence interval

### Logistic regression of SAS score with pattern of smartphone use and temperament

The mean total SAS score of all study subjects is 77.90 (SD 11.73). About 51% of students have above average SAS score. Multivariate analysis was done using logistic regression with three statistically-significant variables: 6 hours or more of smartphone use per day, using smartphone for entertainment, and high HA score (Table 5). Hosmer and Lemeshow test showed that this model is acceptable with p value of 0.722.

**Table 5.** Result of logistic regression multivariate analysis.

| Variable | B | p value | Odds Ratio | 95% CI |  |
| --- | --- | --- | --- | --- | --- |
|  |  |  |  | Lower limit | Upper limit |
| ≥ 6 hours of smartphone use per day | 0.590 | 0.072 | 1.804 | 0.948 | 3.431 |
| Entertainment | 0.998 | 0.102 | 2.713 | 0.820 | 8.977 |
| High HA | 0.710 | 0.020* | 2.035 | 1.119 | 3.701 |
| Constant | -0.735 | 0.023 | 0.479 |  |  |

Hosmer dan Lemeshow test $X^2=2.073$ , $df=4$ , $p=0.722$ ; Nagelkerke $R^2=0.085$ ; $*p<0.05$ .

Through this multivariate model, high HA score is found to be statistically significant (OR 2.035; 95% CI 1.119 to 3.701), while the other two variables are considered as confounding factors: duration of daily smartphone use (OR 1.804; 95% CI 0.948 to 3.431) and using smartphone for entertainment (OR 2.713; 95% CI 0.820 to 8.,977).

## Discussion

This study is the first to report the temperament profile of young adults and relate it with the risk of smartphone addiction. Besides temperament profile, this study also explored other factors such as age, marital status, gender, duration of use, and age at first smartphone use.

Mean duration of smartphone use in this study, 7 hours per day, is higher than other studies. Others have reported a mean daily duration of 3.7 hours (17), or less than 1 hour each day (18). This difference may be accounted for by the difference in sample characteristics. Lin et al. studied engineering students who generally use smartphone less than medical students (17). In the Cho and Lee study, the sample was children who are still under their parent’s supervision (18). This study did not detect a relationship between duration of use and risk of smartphone addiction. However, other studies found such relationship (19,20). One of the diagnostic criteria of addiction is tolerance. A person with addiction will spend more time using smartphones to get the satisfaction.

Younger exposure is also an important factor in the development of addiction. The age at first smartphone use in this study is relatively early, 13 years. Ching et al. proposed that younger age at first use is related to higher risk of addiction (21). Teens are considered vulnerable to addiction as their prefrontal cortex has not matured yet, so they cannot adequately judge the cause and effect of their actions (22,23). A trans-generational study of Zhitomirsky-Geffet and Blau showed that if mobile phone exposure started at teenage years, addiction is easier to happen because their personality is still easily influenced (24).

Activities done using smartphone is quite variable. As a basic need, smartphone is used as a communication device, for calls or short text messages, but nowadays more people use their phones to access social media, browse for information (including shopping), and play games for a long period of time (25). Use of smartphone has shifted from communication to entertainment (26,27).

The search for entertainment is an important theme as it reflects a person’s coping mechanism (28,29). This is supported by Nayak, who proposed that smartphone use in students is related to a wish for escape from reality, even just temporarily (30). Such avoidant coping is often associated with health problems and depression (31). Not only with psychopathology, coping mechanism is related to temperament. The temperament most closely related to avoidant pattern is high harm avoidance (32).

The subjects of this study is homogenous because they come from the same stage of their medical studies. Eley et al. found that medical students are a relatively homogenous group in many aspects, such as age of early 20’s, good educational background, and upper-middle socioeconomic class (9). This study also found other similar characteristics: sex and marital status. This similarity may influence other results such as temperament type and type of smartphone activity.

More study samples are female. They also have a tendency toward HA. Similar to previous studies, temperament is related to gender. Higher HA in female is caused by difference in monoamine system and hormonal composition. With high HA, female has a more active serotonin and noradrenaline components, as well as estrogen that is more sensitive to the HPA axis, making females more vulnerable to anxiety (33). Additionally, female are faster to develop more mature socialization skills compared to male (7).

Female’s tendency to socialize more concurs with the findings of this study: compared to male, females are more likely to use smartphones for social media. Other studies explained females’ use of smartphones for social media by their needs to have wider and deeper relationships, compared to males who tend to only have wider relations (1,11). Through smartphones, females with high HA can socialize more conveniently.

Males use smartphones for online and offline games in this study. Compared to female, male are more likely to be active playing games as they have a more task-oriented motivation (34). Majority of games have this element so they become more interesting for males. Furthermore, games can also become a way to channel males’ aggression (27). Eastern societies, especially Asians, have a higher mean HA scores compared to Americans and Europeans (35). Aggresion tends to be channeled into non-threatening activities, such as games (36), because Asian communities are less individualistic and more communal (35). Besides culture, channeling of aggression in cyberspace is associated with biological factors (brain stucture, hormones, neurotransmitter) that influences social cognition. Smartphone use for social media and games is a good thing up to a certain point, as overuse is associated with avoidant coping, as discussed previously.

Most samples in this study have high HA. Besides gender, this temperament profile can be explained by the sample’s origins in medical faculties. Eley et al. found similar findings in two studies of medical students in Australia: low NS, high RD, and high HA (10). Lee et al. concluded that with high HA, combined with high CO and SD, medical students can develop into independent, reliable, warm, empathic, and friendly doctors (12). However, this type of individuals is also vulnerable to burnout, anxiety, and depression (9,10,37).

In this study, someone with high HA is almost two times more vulnerable to high risk of smartphone addiction. This finding is similar to other studies in internet addiction (38–40). A person with high HA is more likely to be addicted to smartphones compared to students with high NS. This can be explained by the low persistence in NS, making the individual easier to be bored and impulsively shift to other activities.

Duration of smartphone use, 6 hours or more, is not significantly associated, but this variable is a confounding factor and is very important in making a person vulnerable to smartphone addiction. Other studies have also found the relationship between duration of smartphone use and smartphone addiction (13,19,41). This duration also shows a person’s persistence and priorities, as well as the impairment in daily life that may occur. Consequently, change of self priorities is one of the diagnostic criteria of addiction, in DSM 5 and ICD-11 (42,43).

Smartphone use for entertainment show a positive relationship with vulnerability to smartphone addiction. Two studies found a significant relationship between smartphone use for entertainment and smartphone addiction (13,44). Boumosleh and Jaalouk explained that the search for entertainment through smartphones is related to a negative coping mechanism when dealing with depression and anxiety (13). The more frequently a person uses this coping mechanism, the more frequent the use of smartphone for entertainment. The assumption is that this longer duration of use will raise the risk of smartphone addiction, like a vicious cycle; however, this is not studied in this research, requiring further studies.

In general, this study shows that smartphone addiction has similarities with other behavior addictions, such as pathological gambling, internet addiction, and online game addiction (45). Even though not all factors contributing to smartphone addiction are studied, this study shows that there are other factors that may play a role in the vulnerability to smartphone addiction, such as education, social, and personal factors (1,38,40,46).

There are several limitations to this study. First, this study is conducted on only a small proportion of medical students in Jakarta and did not compare between them and non-medical sample, limiting the generalizability of the results. Second, this study did not take into account the types of application used through smartphones, when there is, in fact, some multifunctional application. Additionally, the type of game is not specified, whether online or offline, action, adventure, role play, or puzzles. If the type of application and games are known, further studies can be more specific. Third, there is no consensus on the cutoff score for SAS, so the classification into high and low risk is unclear. Replication of this study with different cutoff will yield different results. Fourth, this study is focused on temperament as a risk factor of smartphone addiction. There are other factors that influence both temperament and smartphone addiction that have not been explored, such as genetics, parenting, character, self-regulation, self-image, comorbidities with other mental disorders, and peer pressure. The impacts of smartphone use in the individual level was not included in this study either.

This study concludes that HA temperament is associated with the risk of smartphone addiction. The higher one’s tendency for HA, the higher the risk of smartphone addiction. Screening on risk of smartphone addiction based on temperament type should be done on medical students. Exposure to behavioral addiction knowledge could be done as early as possible in the medical study as one of the prevention measures of addiction. Further research needs to be conducted on other biological, psychological, and social factors that can also affect a person’s vulnerability to smartphone addiction. Assessment of psychopathology factors on the subject as one of the factors influencing smartphone addiction.

## Acknowledgment

We would like to thank Adhitya Sigit Ramadianto, MD who helped in preparing this manuscript.

## Role of funding source

The research was supported by grant from Universitas Indonesia, named Hibah Publikasi Terindeks Internasional Untuk Tugas Akhir Mahasiswa UI 2018.

## Conflicts of interest

The authors declare that there is no conflict of interest regarding this article.

Authors’ contribution
EH : study concept and design, analysis and interpretation of data, statistical analysis, obtained funding KS : study concept and design, analysis and interpretation of data, statistical analysis, obtained funding, study supervision TW : study concept and design, analysis and interpretation of data, statistical analysis, study supervision IK : study concept and design, analysis and interpretation of data, obtained funding, study supervision MW : obtained funding

